# Prevalence of pathogenic bacteria detected by qPCR from cultured Nile tilapia (*Oreochromis niloticus* Linnaeus, 1758) in southwest Mexico

**DOI:** 10.1101/2023.06.27.546743

**Authors:** Sonia A. Soto-Rodriguez, Francis I. Marrujo Lopez, Karla G. Aguilar-Rendon

## Abstract

Nile tilapia (*Oreochromis niloticus*) is one of the most important aquaculture species in the world. When bacteria are present in cultured tilapia but do not cause a declared disease, it makes them asymptomatic carrier organisms. Once environmental or nutritional conditions change, an outbreak may occur. This is why it is so important to detect pathogens before outbreaks occur. This is the first study that use molecular techniques based on PCR to estimate prevalence of fish pathogens in southwest Mexico. During 2018, 2019, 2020 and 2022 samples of internal organs and lesions of Nile tilapia were taken and analyzed for detection of the main bacterial tilapia pathogens using one-step PCR or qPCR. A total of 2396 samples from the internal organs of Nile tilapia pond and cage cultured come from the Mexican Pacific southwest states of Guerrero, Oaxaca and Chiapas were analyzed. Most of the sampled tilapias were apparently healthy and had no relation between the clinical signs and the pathogens detection was found. No *Francisella* sp. was detected in any sample, *Staphylococcus* sp. was the most prevalent bacterial genus from the three states over time (from 0 to 64 %). Prevalence of *Aeromonas* sp. was from 0 to 4.3 %, although the fish pathogen *A. dhakensis* was not detected. Meanwhile, *S. iniae* was only detected in Chiapas in 2019 at low prevalence (1.4 %) and *S. agalactiae* was detected in the three sites at high prevalence (from 0 to 59 %). Both *Streptococcus* can cause streptococcosis, the most dangerous re-emergent disease to cultured tilapia, which means a great risk for tilapia farming in Mexico.

## INTRODUCTION

Tilapia is the common name of several cichlid species, which are farmed worldwide, with a global annual production that has been growing over time. Nile tilapia (*Oreochromis niloticus*), which is native to northern Africa, currently is the third most abundantly farmed finfish species worldwide, with commercial production recorded in 74 countries. The largest producer country is China (1,241,410 t), followed by Indonesia (1,172,633 t), Egypt (954,154 t), Brazil (343,596 t), and Thailand (205,971 t) (FAO, 2022). The world tilapia aquaculture production grew 10.4 % per year, from around 380,000 tons in 1990 to 6 million tons in 2018 (FAO, 2020a). Thus, tilapia is not only an important food source worldwide but is also one of global economic importance. In general, tilapia is characterized by rapid growth, they are very resistant to low oxygen levels (below 4 mg O_2_ L^-1^), high concentration of organic matter in the water (Arguedas et al., 2017) and is able to survive in high variations of salinity and temperature (Fajer-Ávila et al., 2017). The increased production of farmed Nile tilapia, particularly in tropical rural areas, is primarily because of fish adaptability to diverse production systems, including extensive pond culture, semi-intensive cage culture and intensive methods in ponds, lake cages, tanks, and shrimp ponds (Watanabe et al., 2002). Mexico is currently the ninth largest producer of tilapia in the world, which represents a food solution for the population from southwestern Mexico (CONAPESCA, 2017). However, as in any other farming industry, tilapia is not exempt from diseases associated with ectoparasites, bacteria, virus and fungus, although, most of research papers are focused on parasitic diseases. Cultured tilapia is susceptible to several bacterial diseases caused by pathogens such as *Francisella noatunensis* subsp. *orientalis* (Soto et al., 2013), *Streptococcus agalactiae* (Kayansamruaj et. al., 2014), *Streptococcus iniae* (Anshary et al., 2014); *Edwardsiella tarda* (Clavijo et al., 2002), *Edwardsiella ictaluri* (Soto et al., 2012), and several aeromonads causing Motile Aeromonas septicemia including *Aeromonas sobria* (Li & Cai, 2011), *Aeromonas dhakensis* (Soto-Rodriguez et al., 2018) and *Aeromonas veronii* (Dong et al., 2015). These pathogens are distributed throughout tropical and temperate regions where Nile tilapia are commonly cultured.

Cocci Gram-positive bacteria *S. agalactiae* also known as Group B *Streptococcus* (GBS) and *S. iniae,* a re-emerging bacterial pathogen in freshwater and marine aquaculture, are the causal agents of streptococcosis (Maulu et al., 2021). High prevalence of both pathogens has been observed in cultured tilapia when water temperature was over 27 °C (Mian et al., 2009; Rahmatullah et al., 2017). GBS has been responsible for more than 90 % of the pathogens isolated from farmed dead tilapia each year resulting in a mortality rate of 30 % to 90 % (Liu et al., 2019). Monthly prevalence ranged from 0 to 32 % throughout the year in red hybrid tilapia (Amal et al., 2013). Clinical signs of the acute form of *S. agalactiae* infections include, but are not restricted to erratic swimming, c-shaped body of the fish, exophthalmia (with or without corneal opacity), distended abdomen and hemorrhages (Buller, 2004). So, the economic losses for the tilapia aquaculture industry caused by *S. agalactiae* and *S. iniae* have been huge and, these bacteria are recognized as causative agents of zoonosis (Haenen et al., 2023). Studies on tilapia infected by *Streptococcus* are focused mainly on the isolation, identification and typing of strains, screening of drugs and vaccine development (Cai et al., 2020; De Sousa et al., 2021). Diagnosis of bacterial diseases commonly include clinical signs, wet mount preparations, necropsy, histopathology, and recently molecular techniques. The identification of *Streptococcus* sp. using biochemical profiles are time consuming and difficult because not all bacteria within this genus are included in databases, causing misidentification. Molecular identification using PCR techniques of potential pathogens is the key to early detection of asymptomatic carrier’s fish, unfortunately, rural farmers do not have access to such techniques.

Tilapia from southwestern Mexico (Guerrero, Oaxaca and Chiapas states) is currently cultured in small earthen ponds or floating cages located in dams or small freshwater reservoirs, seeded at low density of fry in semi-intensive systems. Recurrent mortalities, associated with poor management and null implementation of biosecurity measures during cultivation, have been observed over years in these regions. Progress of the cultures are monitored by the aquaculture health committees, however systematic surveillance health programs do not exist. In this context, a monitoring health program was applied focused on the molecular identification of bacterial pathogens in samples of farmed tilapia from mainly rural cultures. In 2018, 2019, 2020 and 2022 samples of internal organs (juvenile and adult) or the entire organism (fry) of tilapia were taken by farmers supported for the aquaculture health committee from each state. Tilapia samples were delivered to the Bacteriology Laboratory to be analyzed using PCR and qPCR techniques. In some years, physicochemical parameters of ponds were registered during collection of the organisms.

## MATERIALS AND METHODS

### Study area

Tilapia farms, consist of rustic earthen ponds or floating cages are in the southwest of the Mexican Pacific Ocean (fig. 1) belonging to the Guerrero, Oaxaca and Chiapas states. From August 2018 to August 2022, samples of Nile tilapia *O. niloticus* (Linnaeus, 1758) were taken to be analyzed.

**Figure 1.**
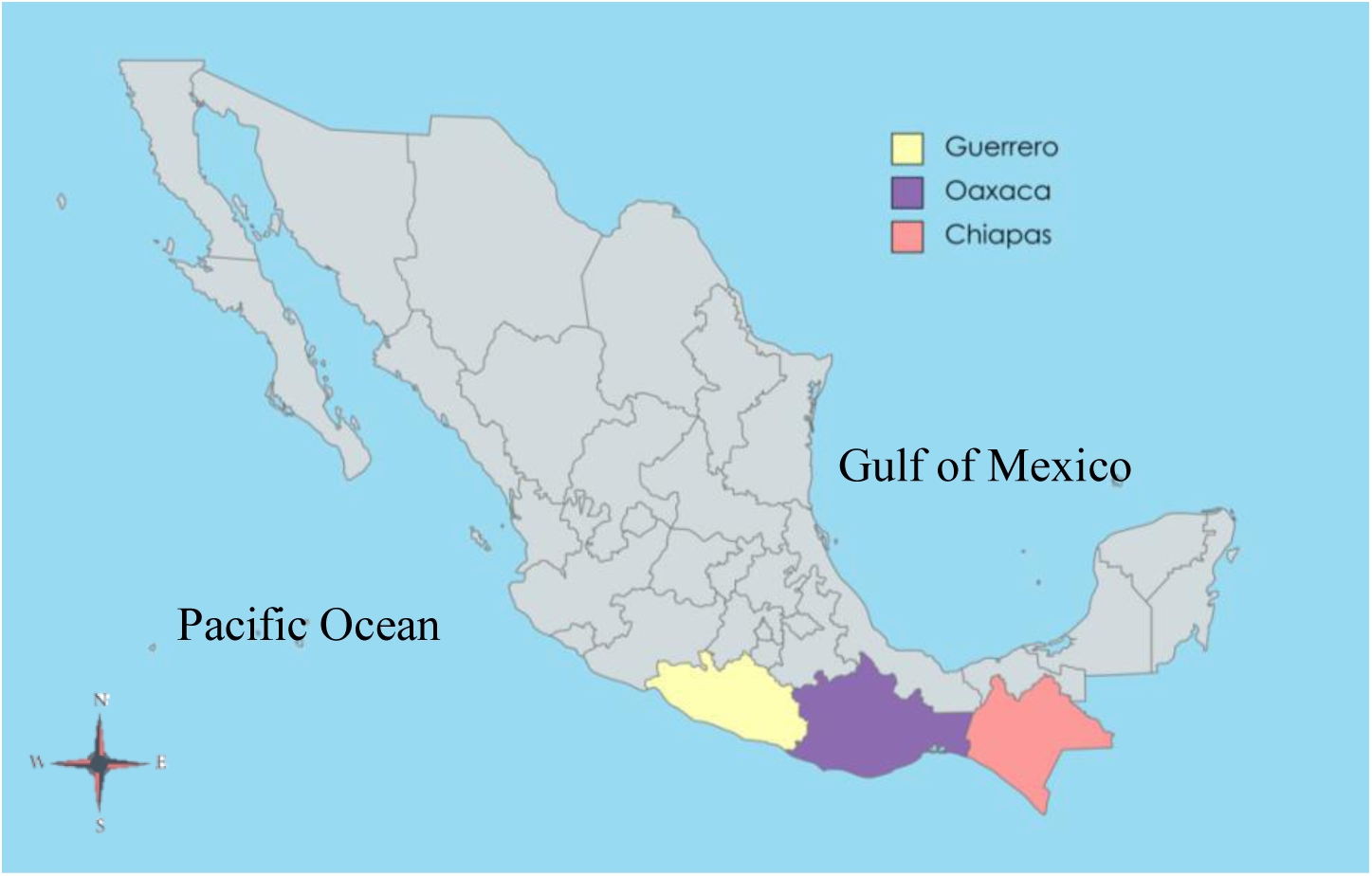
Location of the three southwest Pacific states of Mexico where tilapia (*Oreochromis niloticus*) were collected.

### Collection of organisms

In 2018, 2019 and 2020 field technicians of the State Aquaculture Health Committees from Guerrero, Oaxaca and Chiapas states took samples in each farm, while in 2022 samples were taken by tilapia farmers. Five to six organisms per farm were collected in order to perform analyzes corresponding to the molecular identification of pathogenic bacteria. The number of sampled tilapia farms and samples analyzed is described in Table 1.

**Table 1.** State and number of tilapia tissues analyzed by site and year.

| Year | State | No. farms | No. samples analyzed | Total |
| --- | --- | --- | --- | --- |
| 2018 | Guerrero | 14 | 280 | 760 |
|  | Oaxaca | 12 | 240 |  |
|  | Chiapas | 12 | 240 |  |
| 2019 | Guerrero | 12 | 268 | 800 |
|  | Oaxaca | 12 | 244 |  |
|  | Chiapas | 12 | 288 |  |
| 2020 | Guerrero |  |  | 144 |
|  | Oaxaca | 6 | 144 |  |
|  | Chiapas |  |  |  |
| 2022 | Guerrero | 6 | 148 | 692 |
|  | Oaxaca | 9 | 216 |  |
|  | Chiapas | 14 | 328 |  |
|  | Total | 109 |  | 2396 |

During the sampling, fish were obtained at random directly from the ponds or cages and killed by a puncture in the head to be dissected (OIE 2022). Each tilapia was weighed, measured, and external observations were conducted in each farm. Internally, the aspect, color and contents of the body cavity were also reviewed. Then, samples for bacterial detection from adult and juvenile organisms, were taken from the external skin lesions and the kidney, spleen, liver, and brain were individually dissected. For fry fish, the internal organs were aseptically dissected and polled mixed with the brain. Tissue samples were fixed in ethanol 96° and transported to the Laboratory of Bacteriology at CIAD Mazatlan, Mexico. Water physicochemical parameters were measured in 2018 and 2019 in the culture ponds or floating cages such as oxygen (mg L^-1^), temperature (°C) and pH.

### PCR and qPCR analysis

A total of 2396 samples from tilapia tissues were analyzed. Genomic DNA from samples of kidney, spleen, liver, and brain was extracted using the cetyl trimethylammonium bromide (CTAB) buffer method modified from Doyle and Doyle (1987). The DNA integrity was evaluated by 1 % agarose gel electrophoresis. Genomic DNA from the bacterial strains used as positive controls was extracted using the Wizard® Genomic DNA Purification Kit (Promega, Madison, WI, U.S.) according to the manufacturer’s instructions. The obtained DNA was stored at -20 °C until PCR analysis DNA was quantified using a DeNovix DS-11 Series (Thermo Fisher NanoDrop™ One Spectrophotometer). DNA was analyzed for one-step or real-time PCR (qPCR) amplifications for the pathogens: *Francisella* sp*., Staphylococcus* sp., *Staphylococcu*s *aureus, Staphylococcus gallinarum, Aeromonas* sp., *Aeromonas dhakensis*, *Streptococcus iniae*, and *Streptococcus agalactiae* (Table 2). Strains used as positive control were stored at -80 °C until used.

**Table 2.**
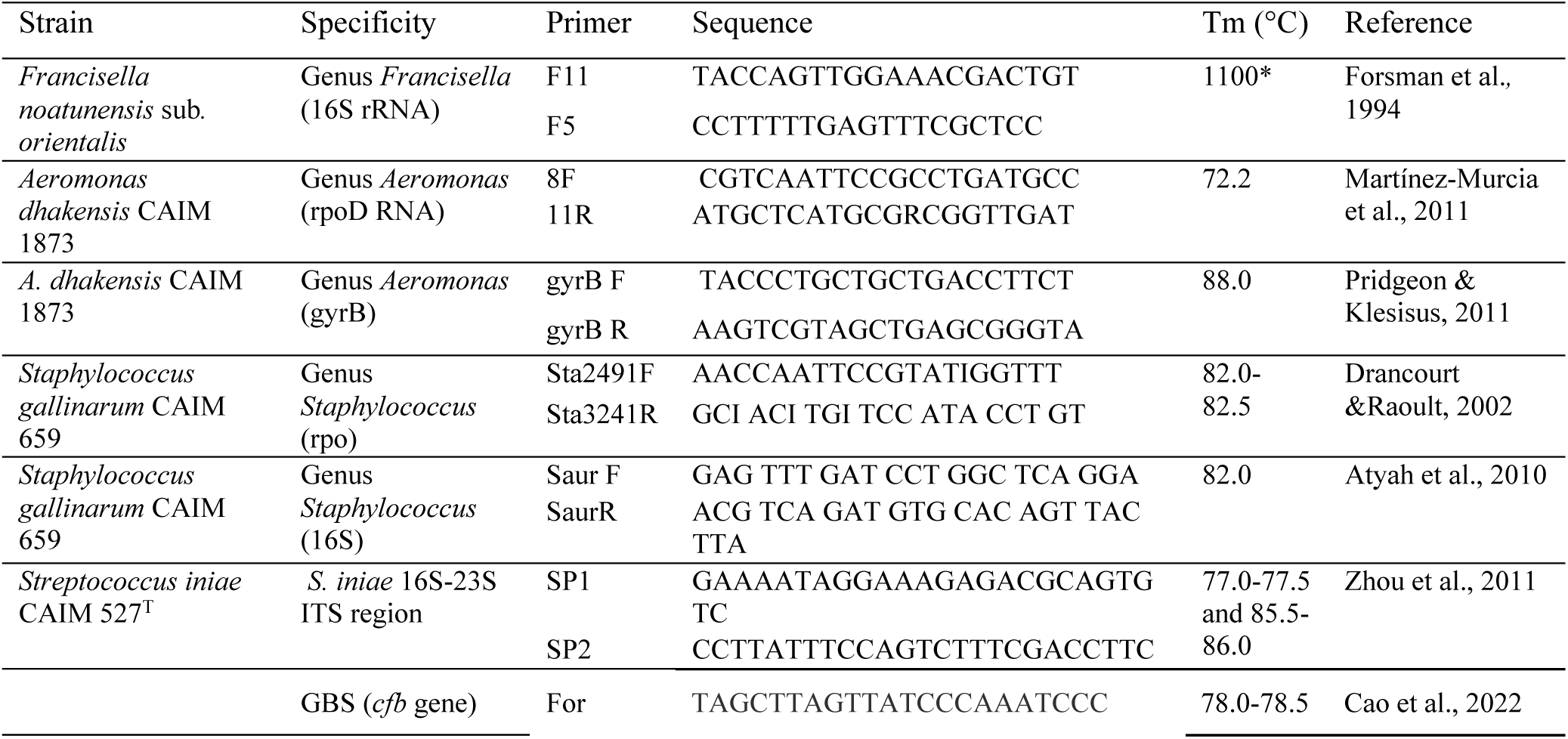
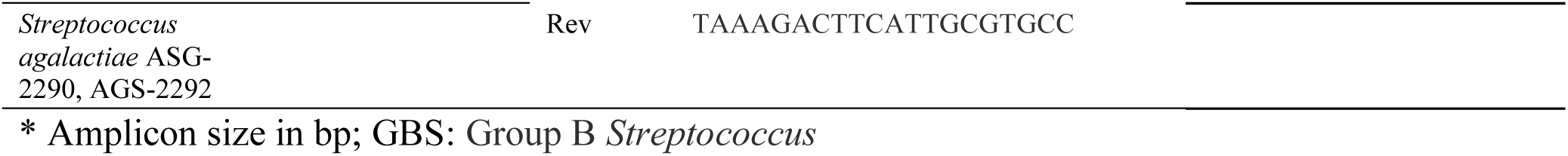
Bacterial strains used as positive control for detection of pathogens to cultured tilapia.

### Detection of *Francisella* using one-step PCR

To confirm the *Francisella* genus, amplification was carried out in a total reaction volume of 11.2 μL, composed of 3.0 μL PCR Master Mix (Thermo Scientific™), 6.0 μL 18 Ohms deionized water, 0.6 μL (0.1 mM) of each primer set (Table 2), and 1.0 μL (25 ng µL^-1^) of DNA template. Amplification was performed with the SimpliAmp Thermal Cycler (Applied Biosystems®) using the following settings: initial denaturalization at 94 °C for 30 s; followed by 35 cycles at, 94 for 3 min, 94 °C for 30 s, 60 °C for 1 min, and 72 °C for1 min; with a final extension at 72 °C for 2 min. Genomic DNA from a *F. noatunensis* subsp. *orientalis* strain, kindly donated by Dr. Esteban Soto (The University of California) was included as a positive control for each PCR assay. Negative control consisted of the same reaction mixture but without DNA. PCR amplification products were electrophoresed in 1.0 % agarose gel at 120 V for 30 min at room temperature. A 100-3000 bp DNA ladder (AXYGEN) was used as a molecular marker. Gels were visualized under Axygen® Gel Documentation Systems (Corning).

### Detection of bacterial pathogens using qPCR

For detection of *Staphylococcus* sp., *S. aureus*, *S. gallinarum, Aeromonas* sp., *A. dhakensis*, *S. iniae* and, *S. agalactiae* amplification was carried out in a total reaction volume of 5.0 μL, composed of 2.5 μL Luna® Universal qPCR Master Mix (New England Biolabs, UK), 1.8 μL 18 Ohms deionized water, 0.1 μL (0.1 mM) of each primer set, and 0.5 μL (25 ng µL^-1^) of DNA template. Strains used as positive control and sets of primers are in Table 2. Negative control consisted of the same reaction mixture but without DNA. All qPCR amplification reactions were carried out using a CFX96 Touch Real-Time PCR Detection System (BioRad), a two-positive control (a sample including genomic DNA from strain bacterial) and two-negative control (a sample without genomic DNA) were added to each reaction. The PCR protocol for *Staphylococcus* sp. and *S. aureus* consisted of 1 cycle of 30 s at 94 °C, followed by 35 cycles of 30 s at 94 °C, 30 s at 55 °C and 1 min at 72 °C and final extension from 65 at 95 °C every 5 s at 5 °C. The PCR scheme for *Aeromonas* sp. consisted of 1 cycle of 30 s at 94 °C, followed by 35 cycles of 30 s at 94 °C, 30 s at 55 °C and 1 min at 72 °C and final extension from 65 at 95 °C every 5 s at 5 °C. The PCR scheme for *S. iniae* consisted of 1 cycle of 30 s at 94 °C, followed by 35 cycles of 30 s at 94 °C, 1 min at 60 °C and 1 min at 72 °C and final extension from 65 at 95 °C every 5 s at 5 °C. The PCR protocol for *S. agalactiae* consisted of 1 cycle of 3 min at 94 °C, followed by 35 cycles of 15 s at 95 °C, 30 s at 56.6 °C and final extension from 65 at 95 °C every 5 s at 5 °C. After the amplification of each pathogenic, the melting step was performed to evaluate the Tm.

Prevalence of pathogenic bacteria over time and origin was estimated using the online program Win Epi (Working in Epidemiology http://www.winepi.net/uk/index.htm) with 95% confidence level.

## RESULTS

### Clinical signs of collected tilapias

In 2018, most of sampled tilapias from Guerrero, Oaxaca and Chiapas were apparently healthy, some organisms showed eroded fins, white cysts on fins, injured eyes, swollen kidney, liver, and spleen, exophthalmia, friable liver, ulcers, external lesions and desquamation (fig. 2). During 2019 in Guerrero and Oaxaca most organisms were apparently healthy, some tilapias displayed hemorrhagic areas, pale and swollen liver, exophthalmia, swollen spleen, and petechiae in the operculum. However, half of tilapias from Chiapas showed pale liver and few organisms with swollen liver and desquamation. In 2020 there is no data on clinical signs. In 2022 the clinical signs recorded from some organisms presented eroded fins, swollen abdomen, injured skin, exophthalmia, and desquamation.

**Figure 2.**
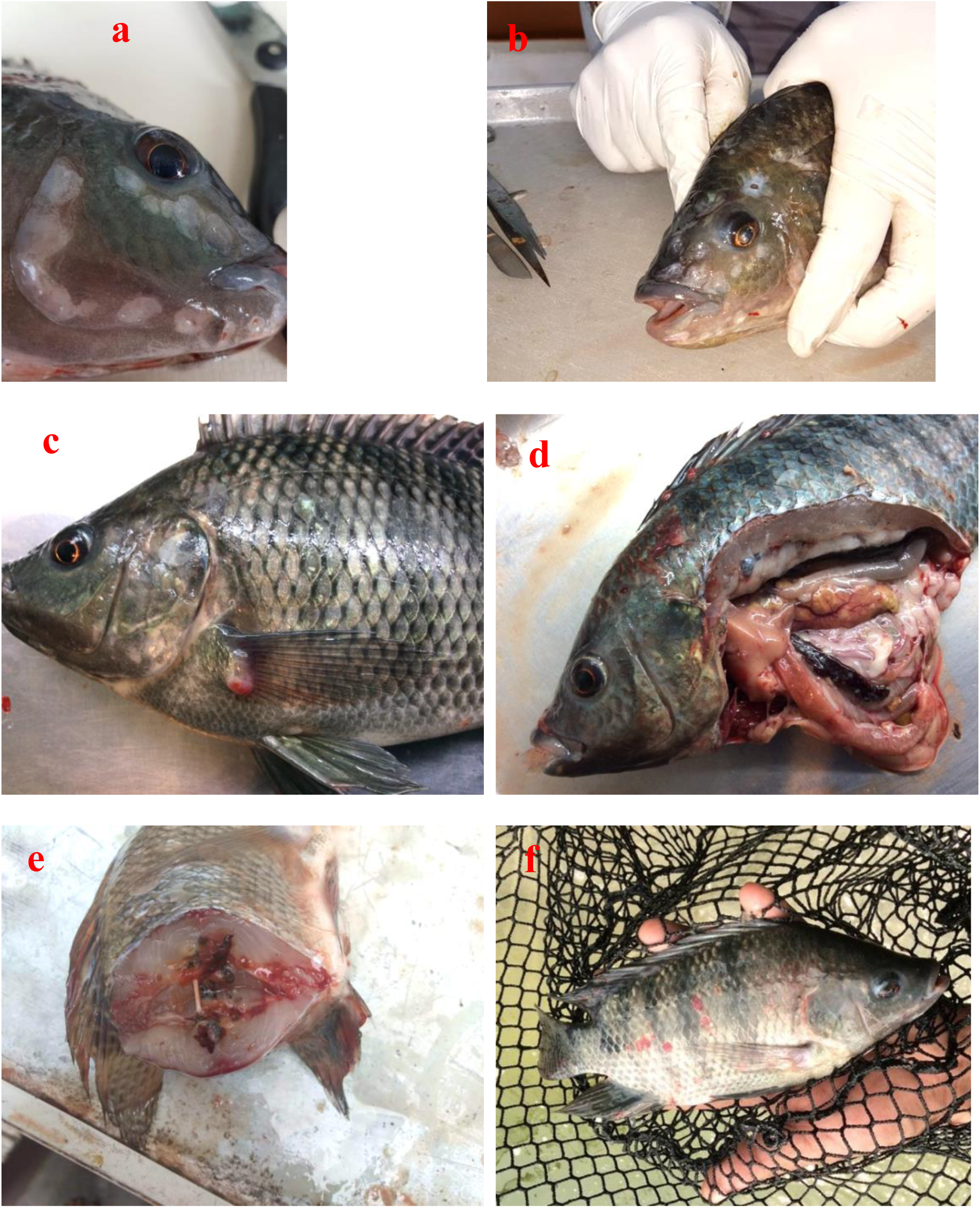
External and internal lesions recorded on farmed Nile tilapia (*Oreochromis niloticus*) from Guerrero, Oaxaca and Chiapas. **a**, ulcers on mandibula; **b**, lesions on the head; **c**, cyst on pectoral fin; **d**, pale and swollen liver, white granule in spleen and cystson skin; **e**, hemorrhage and necrotic lesions on skeletal muscle; **f**, hemorrhage on skin and opercula.

### Physicochemical parameters

In 2018 the water temperature ranged from 21 to 32 °C, while in 2019 it was slightly higher from 24 to 33 °C (see Table 1 in Supplementary Material). Dissolved oxygen in 2019 was between 2.8 and 11.0 mg L^-1^, while 2019 was from 2.3 to 13.0 mg L^-1^. Values of pH were more homogeneous during the sampling time ranging from 7.0 to 9.0 during both years.

### Detection of bacteria

No *Francisella* genus was detected in all samples of tilapia from Guerrero, Oaxaca and Chiapas in the time period sampled using the one step PCR technique (see fig. S1 in Supplementary Material). Fig S1 shows the amplicon (1100 bp) of the positive control for detection of *Francisella* sp. and negative tissue samples of the internal organs without amplification from Chiapas collected in 2019. While *Staphylococcus* genus was the most detected bacteria from the three states over time (fig. 3). The pathogen *S. iniae* was only detected in Chiapas in 2019 at low prevalence (1.4 %) and *S. agalactiae* had the highest prevalence (59 %) detected in Guerrero in 2022.

**Figure 3.**
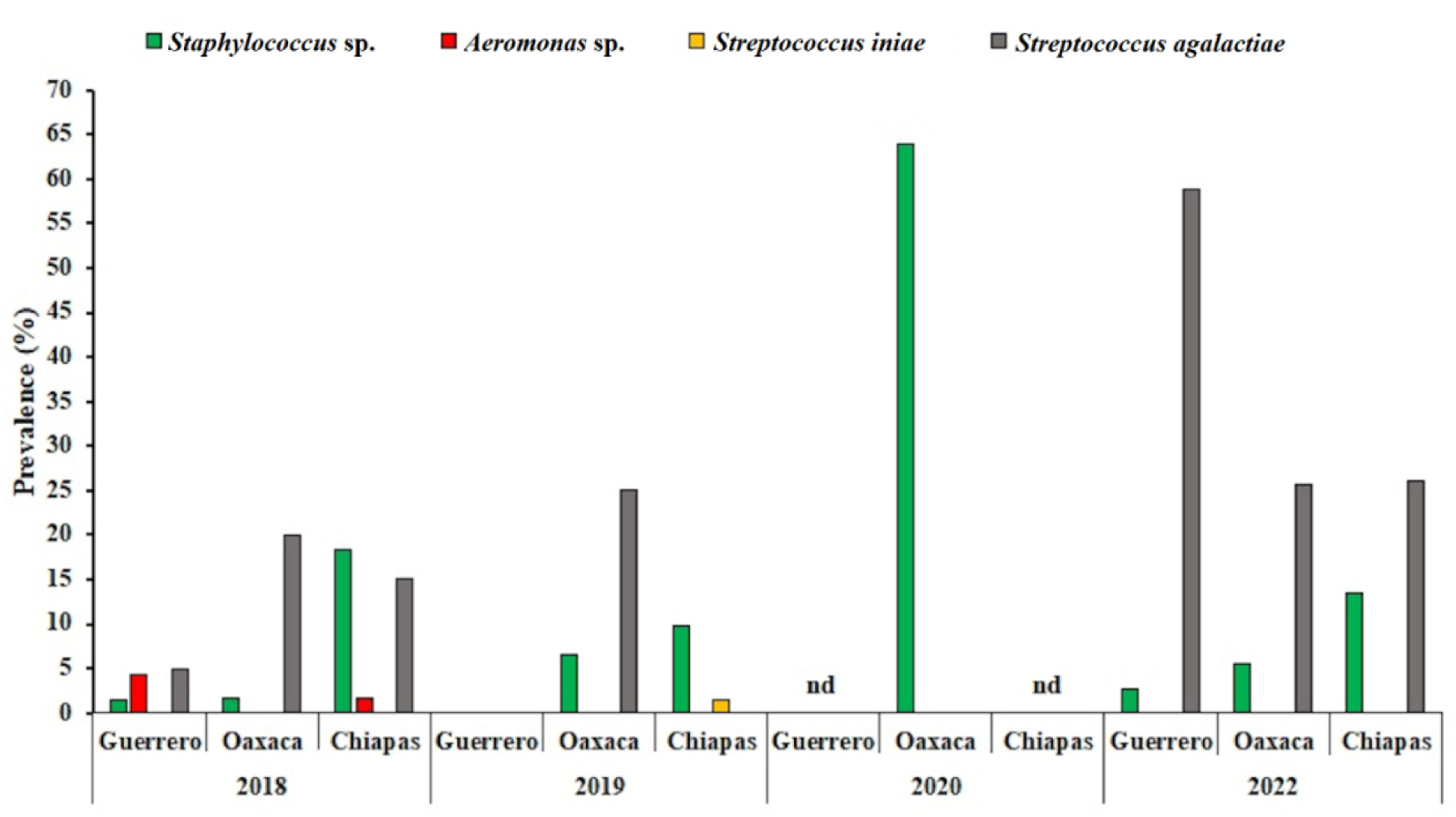
Percentage of prevalence of pathogenic bacteria of farmed Nile tilapia (*O. niloticus*) over time in Guerrero, Oaxaca and Chiapas southwest Mexico. nd, no sample.

*Aeromonas* sp. was detected at low prevalence in 2018 in samples collected from Guerrero (4.3 %) and Chiapas (1.7 %), however *A. dhakensis* was not detected from any internal organs sample of tilapia (fig. 4).

**Figure 4.**
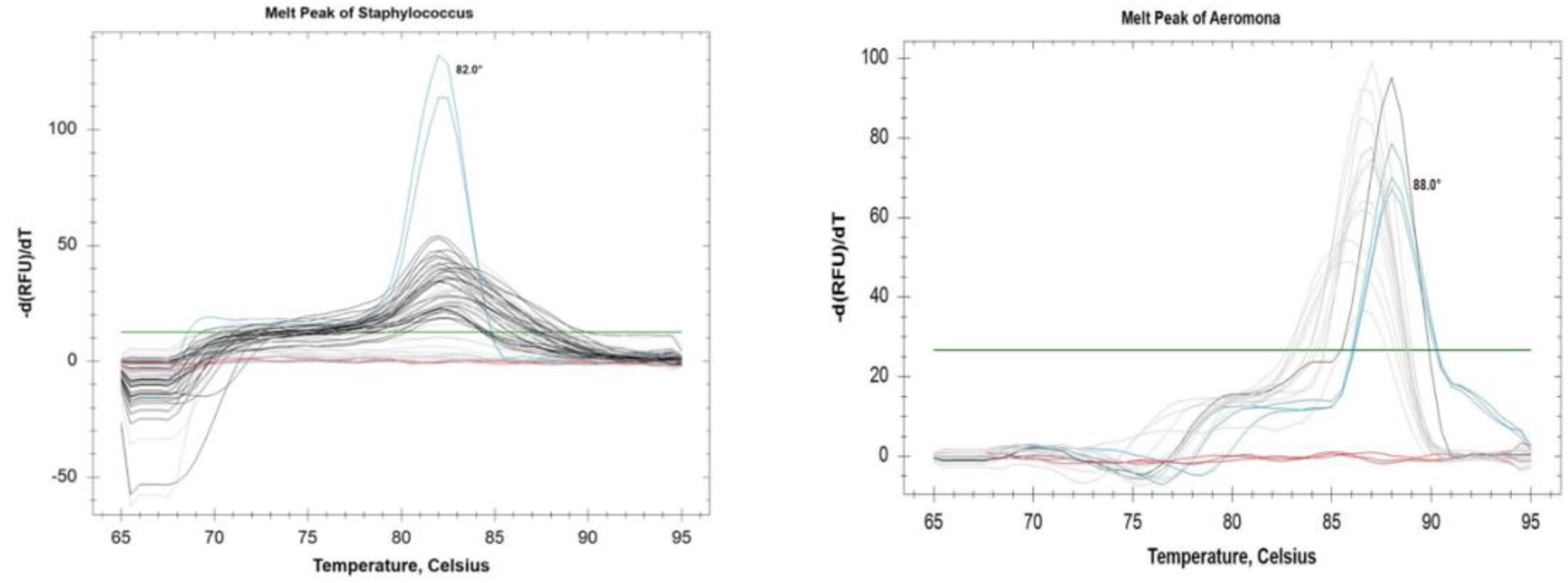
Positive control (Tm 82.0 °C) of *Staphylococcus gallinarum* CAIM 659 (blue line), examples of positive detection (black lines) and negative detection (gray lines) for *Staphylococcus* sp. in tissue samples of *O. niloticus* from Oaxaca and Chiapas collected in 2022. Positive control (Tm 88.0 °C) of *Aeromonas dhakensis* CAIM 1873 (blue line), examples of positive detection (black line) and negative detection (gray lines) for *Aeromonas* sp. in tissue samples of *O. niloticus* from Chiapas collected in 2018. Negative controls (red lines).

*S. iniae* was only detected at low prevalence (1.4 %) in Chiapas in 2019. Positive control (*Streptococcus iniae* CAIM 527^T^) showed two Tm, first at 77.0-77.5 °C, the second at 85.5-86.0 °C (fig. 5). The highly pathogenic Nile tilapia *S. agalactiae* was detected at different prevalence’s in 2018, 2019 and 2022. The highest prevalence was in 2022 in Guerrero (59 %) (fig. 3). The Tm of the positive control (*S. agalactiae* ASG-2290) was 78.0-78.5 °C (fig. 5).

**Figure 5.**
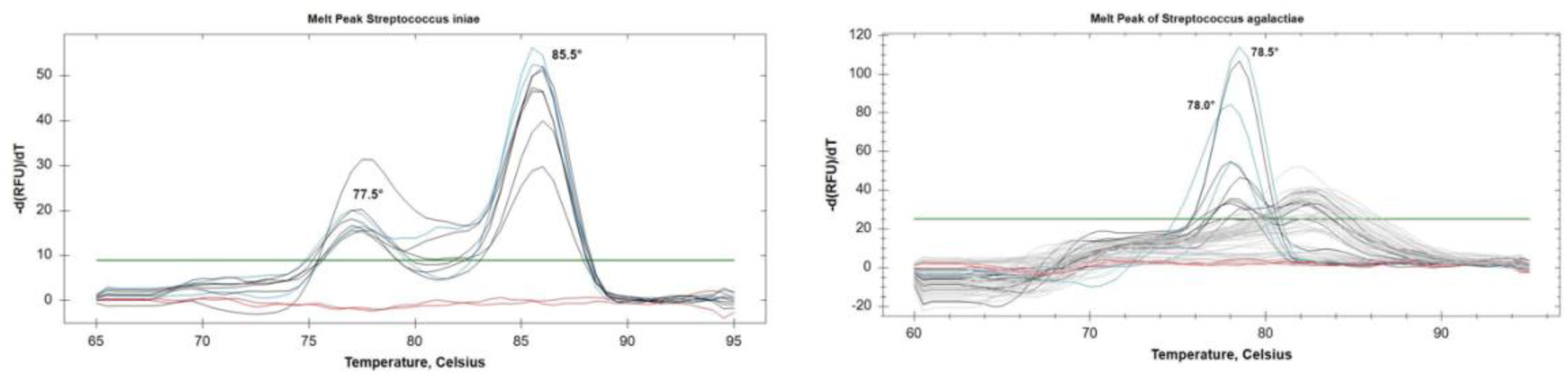
Positive control (77.5° and 85.5 °C) for *Streptococcus iniae* CAIM 527^T^ (blue lines), examples of positive detection (black lines) and negative detection (gray lines) for *S. iniae* in tissue samples of *O. niloticus* from Chiapas collected in 2019. Positive control of *Streptococcus agalactiae* ASG-2290 (78.0-78.5 °C) (blue lines), examples of positive detection (black lines) and negative detection (gray lines) for *S. agalactiae* in tissue samples of *O. niloticus* from Guerrero collected in 2022. Negative controls (red lines).

## DISCUSSION

The Pacific white shrimp (*Penaneus vannamei*) and tilapia were the two largest species with 64 and 21 % respectively of the total production (FAO, 2020a), and Mexico is the fourth-largest aquaculture country in Latin America and the Caribbean (FAO, 2020b). Rural production in Mexico has a significant contribution to the food security and livelihood of numerous households and contributes to improving family nutrition, generating extra income, keeping family intact, discouraging emigration, and empowering women within the family (Martínez-Cordero et al., 2021). The Pacific southwest Mexico states (Guerrero, Oaxaca and Chiapas) have traditionally lived in poverty; in 2020 over 40 % population belongs to the rural people and 87 % of the adult people was illiterate (INEGI, 2023). Furthermore, the rural people that work in the tilapia culture speak one or more indigenous languages, which makes communication difficult. The above implies greater difficulty in accessing training courses and workshops on tilapia farming. These communities are supported by the State Aquaculture Health Committees of each state, but the long distance among farms makes it difficult this task during infectious outbreaks. In addition, farmers do not keep records of mortality events, commercial products or treatments that they use during the culture.

In this multi-year study (2018, 2019, 2020 and 2022), several tilapia farms were sampled to detect fish pathogens. New emerging or re-emergent bacterial fish diseases, rose as a new threat for aquaculture, which may be attributed to close contact of the aquaculture environment with animal and/or human wastes. From 2018 to 2020, most of the sampled tilapias from Guerrero, Oaxaca and Chiapas were apparently healthy. However, the swollen livers and spleens were the most frequent clinical signs, which were also observed in apparently healthy tilapias cage cultured in damps from Sinaloa state, located in northwestern Mexico and no pathologies were associated with the clinical signs (Soto-Rodriguez et al., 2013). In this study, in 2022, most of data were missed, although, some organisms displayed swollen abdomen resulting from ascites, injured skin, exophthalmia, and desquamation, signs associated with *Streptococcus* infection in Nile tilapia (Amal & Zamri-Saad, 2011).

*Staphylococcus* was present in the three states over time with the highest prevalence followed by *S. agalactiae*. *S. aureus* was reported in *O. niloticus* triggering high mortality with various pathological alterations (Gaafar et al., 2015). It also poses health risks to fish handlers and consumers (Grema et al., 2015). Methicillin resistant *S. aureus* has been related to mortality in the culture of Nile tilapia in northern Egypt (Soliman et al., 2014) and Malaysia (Atyah et al., 2010). Naturally infected *O. niloticus* with *S. aureus* showed hemorrhages on the skin and erosions at the dorsal region, swollen congested kidney, congested gills and pale liver (Soliman et al., 2014). Pathological changes ranged between minor degenerative modifications to severe necrotic alterations mainly in the brain, kidney, spleen and gills (Gaafar et al., 2015). Although in this study *S. aureus* was not detected, it would be worthwhile to seek new molecular methodologies for its detection. Motile Aeromonas septicemia comprise several species which posse lipases, aerolysins, serine protease, cytotoxic enterotoxins, hemolysins virulence factors (Soto-Rodriguez et al., 2018; El-Gohary et al., 2020). Although the etiological agent of this disease has been usually reported as *A. hydrophila* most of the *Aeromonas* species has been incorrectly identified (Soto-Rodriguez et al., 2018). In this study, *Aeromonas* sp. was only detected at low prevalence in 2018 in Guerrero and Chiapas, but not the fish pathogen *A. dhakensis*. Soto-Rodriguez et al. (2009) reported 30 % prevalence of presumably *A. hydrophila* (identified using biochemical protocols) in cage cultured tilapia from Sinaloa damps. Bacteria was isolated from the liver, kidney and brain of tilapias displaying desquamation, blindness, protruding and opaque eyes, eroded caudal fin and pale gills. In this study, *S. iniae* was only detected at low prevalence in 2019 in Chiapas, the leading producer of farmed tilapia in Mexico. While, *S. agalactiae* was found at high prevalence in 2022 in Guerrero, Oaxaca and Chiapas (59, 26 and 26 % respectively), which constitutes a threat to the state’s aquaculture. Differences in detection of pathogens among three states over time could be due to the selected farms (size, production system, distance, logistics, implementation of good practices, among others). *Streptococcus* was presumably detected in 2009 in the liver and spleen of apparently healthy cage cultured tilapia from Sinaloa damps (Soto-Rodriguez, 2009). *S. agalactiae* may cause acute or chronic disease in tilapia. An outbreak or cumulative mortality during chronic phase in tilapia can reach 80 % (FAO, 2021). In the chronic form, light brown or dark red nodules were seen in the skeletal muscle near the vertebra of Nile tilapia (Li et al., 2014), lesions also observed in this study (fig. 2e). In addition, this disease is highly contagious, easily transmitted from fish to fish, by contact with individuals carrying the bacteria or through water. The disease is directly related to high temperatures and nitrogenous wastes in the water that cause stress to the fish, increasing their incidence and vulnerability to the disease, since the immune response of the host to the pathogen is affected.

There are informal reports (published by private corporations who are dedicated to fish vaccination) of *S. agalactiae* isolated from farmed tilapia in Mexico. The Fish Health Forum (sponsored by PharmaQ) reported outbreaks in 2019 caused by *S. agalactiae* serotype 1a, causing tilapia mortality over 60 % in vaccinated batches and up to 85 % in unvaccinated, specifically in hot periods of the year when the water temperature was over 32 °C (https://fishhealthforum.com/emerging-diseases-in-tilapia-farming-in-latin-america/). Recently, the Mexican government reported high prevalence of *S. iniae* in Queretaro (29 %) and Guerrero (33 %) and *S. agalactiae* in Oaxaca (33 %), Queretaro (12 %) and Veracruz (20 %) in cultured tilapia and catfish (SAGARPA, 2022), unfortunately, the method used for bacterial identification was not mentioned. Amplification by PCR and qPCR of 16S rRNA and 16S-23S rRNA intergenic spacer genes can be used to identify distinct *Streptococcus* species (Lau et al., 2003). Ortega et al. (2017) identify *S. iniae* by PCR and 16S rRNA sequencing as the causative disease agent in two tilapia (*Oreochromis aureus*) Mexican populations. Furthermore, *S. agalactiae* and *S. iniae* were identified in 2017 and *Streptococcus* sp. in 2019, using 16S rRNA sequencing from cultured tilapias from a Guerrero dam (Castro Ortiz, 2020). The main clinical signs were erratic swimming, lethargy, exophthalmia, ulcers, desquamation, fin erosion, ascites and friable organs. Similar sings have been recorded for *Enterococcus* sp. and bacterial hemorrhagic septicemia, so timely and accurate diagnosis is a priority. At Malpaso dam located in Chiapas, 80 % of the farms, reaching 50 % mortality, were presumably positive for *Streptococcus* spp., using biochemical methods for identification (Hernández Hernández, 2021). Temperature and oxygen, along with rainy and dry periods, are considered the main climate-related risks in floating cages, as high temperatures predispose stress and susceptibility to disease (Cochrane, 2009; Lebel, 2016). In this study during 2018 and 2019 water temperatures of the tilapia cultures were within the acceptable to optimal levels to grow Nile tilapia (El-Shafai et al., 2004; El-Sayed, 2006) but was not possible to establish a clear relation among the physicochemical variables and pathogens detection due to lack of data. In Mexico, rural tilapia farmers register mortality every year but information about infectious diseases is scarce; the cause, the etiological agent or the predisposing condition is unknown, attributing mortalities to poor water quality and inadequate culture management. In addition, there a lack of appropriate diagnostic tools in these regions.

## CONCLUSIONS

In this study, and high prevalence of a major fish pathogen *S. agalactiae* was found in samples of the internal organs of pond and cage farmed tilapia from the three sampled states (Guerrero, Oaxaca and Chiapas), which is a major threat to sustainable Nile tilapia farming in Mexico. In addition, *Aeromonas* sp. and *S. iniae* were also detected over years and *Staphyllococcus* at high prevalence, whose species might have pathogenic strains, although no evident clinical signs were registered.

## ACKNOWLEDGEMENTS

The authors would like to thank Committee of Aquaculture Health of the State of Guerrero, Oaxacan Committee of Aquaculture Health and Safety, the State Committee for Aquaculture Health of Chiapas for your help during collection of organisms. Thanks to Circe Romero for logistical support on rural farms.

## DATA AVAILABILITY STATEMENT

The manuscript does not contain shared data.

## FUNDING INFORMATION

The present study was supported by CONAHCYT grant FORDECyT 292474 “ Desarrollo regional sustentable en los estados del Pacífico Sur Mexicano: modelo de política pública para la sustentabilidad alimentaria enfocado en el cultivo de tilapia en la micro, pequeña y mediana MIPYME empresa” (Sustainable regional development in the South Pacific states of Mexico: a public policy model for food sustainability focused on tilapia farming in micro, small and medium-sized MIPYME enterprises).

## CONFLICT OF INTEREST

The authors declare that they have no conflict of interest.

## Notes

### Competing Interest Statement

The authors have declared no competing interest.

## REFERENCES

Amal, M. N. & Zamri-Saad, M. (2011). Streptococcosis in tilapia (*Oreochromis niloticus*): a review. Pertanika Journal of Tropical Agricultural Science, 34 (2) 195–206

Amal, M. N., Zamri-Saad, M., Siti-Zahrah, A. & Zulkafli, A. R. (2013). Transmission of *Streptococcus agalactiae* from a hatchery into a newly established red hybrid tilapia, *Oreochromis niloticus* (L.) x *Oreochromis mossambicus* (Peters), farm. Journal of Fish Diseases, 36(8), 735–739. https://doi.10.1111/jfd. 12056

Anshary, H., Kurniawan, R. A., Sriwulan, S., Ramli, R., & Baxa, D. V. (2014). Isolation and molecular identification of the etiological agents of streptococcosis in Nile tilapia (*Oreochromis niloticus*) cultured in net cages in Lake Sentani, Papua, Indonesia. SpringerPlus, 3(1), 627. https://doi.10.1186/2193-1801-3-627

Arguedas D., Ortega C., Martínez S. & Astroza A. (2017). Parasites of Nile tilapia larvae *Oreochromis niloticus* (Pisces: *Cichlidae*) in concrete ponds in Guanacaste, Northern Costa Rica. UNED Research Journal, 9(2), 313. https://doi.10.22458/urj.v9i2.1904

Atyah, M., Zamri-Saad, M. & Siti-Zahrah, A. (2010). First report of methicillin-resistant *Staphylococcus aureus* from cage-cultured tilapia (*Oreochromis niloticus*). Veterinary Microbiol, 144, 502–504. https://doi.org/10.1016/j.vetmic.2010.02.004

Buller, Nicky B. (2004). Buller, N. B. (2004). Bacteria from fish and other aquatic animals: A practical Identification Manual. CABI publishing

Cai, Y. Z., Liu, Z. G., Lu, M. X., Ke, X. L., Zhang, D. F., Gao, F. Y., Cao, J. M., Wang, M. & Yi, M. M. (2020). Oral immunization with surface immunogenic protein from *Streptococcus agalactiae* expressed in *Lactococcus lactis* induces protective immune responses of tilapia (*Oreochromis niloticus*). Aquaculture Reports, 18, 100538. https://doi.org/10.1016/j.aqrep.2020.100538

Cao, J., Liu, Z., Zhang, D., Guo, F., Gao, F., Wang, M., Yi, M. & Lu, M. (2022). Distribution and localization of *Streptococcus agalactiae* in different tissues of artificially infected tilapia (*Oreochromis niloticus*). Aquaculture, 546, 737370. https://doi.org/10.1016/j.aquaculture.2021.737370

Castro Ortiz, A. (2020). Identificación y caracterización de patógenos bacterianos aislados de tilapias del Nilo (*Oreochromis niloticus*) cultivadas en la presa de El Gallo, Guerrero, México. Master Thesis. Universidad Michoacana de San nicolás de Hidalgo, Facultad de Biología, 54 pp http://bibliotecavirtual.dgb.umich.mx:8083/xmlui/handle/DGB_UMICH/2834.

Clavijo, A., Conroy, G., Conroy, D., Santander, J. & Aponte, F. (2002). First report of *Edwardsiella tarda* from tilapias in Venezuela. Bulletin of European Association of Fish Pathologists, 22(4), 280– 282

Cochrane, K., De Young, C., Soto & D., Bahri, T. (2009). Climate change implications for fisheries and aquaculture. FAO. Fisheries and Aquaculture Technical Document, 530, 212 pp.

CONAPESCA 2017. Mexico among the top ten places in tilapia production in the World. https://www.gob.mx/conapesca/articulos/mexico-entre-los-diez-primeros-lugares-en-produccion-de-tilapia-enmundo?idiom=es#:~:text=La%20importancia%20de%20la%20tilapia,totalidad%20es%20por%20sistemas%20acu%C3%ADcolas. [Accessed 15 March 2023]

De Sousa, E. L., Assane, I. M., Santos-Filho, N. A., Cilli, E. M., De Jesus, R. B & Pilarski, F. (2021). Haematological, biochemical and immunological biomarkers, antibacterial activity, and survival in Nile tilapia *Oreochromis niloticus* after treatment using antimicrobial peptide LL-37 against *Streptococcus agalactiae*. Aquaculture, 533, 736181. https://doi.org/10.1016/j.aquaculture.2020.736181

Drancourt, M., & Raoult, D. (2002). rpoB gene sequence-based identification of *Staphylococcus* species. Journal of Clinical Microbiology, 40(4), 1333–1338. https://doi.10.1128/JCM.40.4.1333-1338.2002

Dong, H. T., Nguyen, V. V., Le, H. D., Sangsuriya, P., Jitrakorn, S., Saksmerprome, V., Senapin, S. & Rodkhum, C. (2015). Naturally concurrent infections of bacterial and viral pathogens in disease outbreaks in cultured Nile tilapia (*Oreochromis niloticus*) farms. Aquaculture, 448, 427–435. https://doi.org/10.1016/j.aquaculture.2015.06.027

Doyle, J. J. & Doyle, J. L. (1987). A rapid DNA isolation procedure for small quantities of fresh leaf tissue. https://worldveg.tind.io/record/33886/

El-Gohary, F.A., Zahran, E., Abd El-Gawad, E.A., El-Gohary, A.H. M. Abdelhamid, F., El-Mleeh, A., Elmahallawy, E.K. & Elsayed, M.M. (2020). Investigation of the prevalence, virulence genes, and antibiogram of motile aeromonads isolated from Nile tilapia fish farms in Egypt and assessment of their water quality. Animals, 10, 1432. https://doi.org/10.3390/ani10081432

El-Sayed, Abdel F. M. (2006) Tilapia Culture. Oxfordshire: CABI Publishing. Wallingford, UK, 277 pp https://doi.org/10.1079/9780851990149.0000

El-Shafai, S.A., El-Gohary, F.A. & Nasr, F.A. (2004). Chronic ammonia toxicity to duckweed-fed tilapia (*Oreochromis niloticus*). Aquaculture 232, 117–127. https://doi.org/10.1016/S0044-8486(03)00516-7

Fajer-Ávila, E. J., Medina-Guerrero, R. M. & Morales-Serna, F. N. (2017) Strategies for prevention and control of parasite diseases in cultured tilapia. Acta Agrícola Pecuaria, 3, 25–31

FAO (2020a). Top 10 species groups in global, regional and national aquaculture 2018. Rome. 316 pp. www.fao.org/3/ca9383en/ca9383en.pdf [Accessed April 2023]

FAO (2020b). Aquaculture growth potential in Latin America and the Caribbean (LAC). Rome. 83 pp. www.fao.org/3/ca8180en/ca8180en.pdf

FAO (2021). Risk Profile-Group B Streptococcus (GBS)-*Streptococcus agalactiae* Sequence Type (ST) 283 in Freshwater Fish https://www.fao.org/publications/card/fr/c/CB5067EN [Accessed April 2023]

FAO (2022). FishStatJ - Software for Fishery and Aquaculture Statistical Time Series. https://www.fao.org/fishery/en/topic/166235?lang=en [Accessed April 2023]

Forsman, M., Sandstrom, G. & Sjöstedt, A. (1994). Analysis of 16S Ribosomal DNA sequences of Francisella strains and utilization for determination of the phylogeny of the genus and for identification of strains by PCR. International Journal of Systematic Bacteriology, 44, 38–46. https://doi.org/10.1099/00207713-44-1-38

Gaafar, A., Soliman, M., Ellakany, H., Affr, N., Elbialy, A., Mona, S.Z., Younes, A. & Abozahra, R. (2015). Comparative pathogenicity of methicillin-resistant *Staphylococcus aureus* (MRSA) in Nile tilapia (*Oreochromis niloticus*) and *Tilapia zilli*. Life Science Journal, 12(3), 186–194

Grema, H. A., Geidam, Y. A., Gadzama, G. B., Ameh, J. A. & Suleiman, A. (2015). Methicillin resistant *Staphylococcus aureus* (MRSA): a review. Advances Animal and Veterinary Sciences, 3, 79–98. https://doi.org/10.14737/journal.aavs/2015/3.2.79.98

Haenen, O. L., Dong, H. T., Hoai, T. D., Crumlish, M., Karunasagar, I., Barkham, T., Chen, S. L., Zadoks, R., Kiermeier, A., Wang, B., Gamarro, E. G., Takeuchi, M., Amal, M. N., Fouz, B., Pakingking, R., Wei, Z. W. & Bondad-Reantaso, M. G. (2023). Bacterial diseases of tilapia, their zoonotic potential and risk of antimicrobial resistance. Reviews in Aquaculture, 15, 154–185. https://doi.org/10.1111/raq.12743

Hernández Hernández, M., Gutiérrez Jiménez, J., Coutiño Estrada, B., Ruiz Sesma, B., & Bautista Trujillo, G. (2021). Oxígeno-temperatura en la incidencia de *Streptococcus* sp. en jaulas flotantes de tilapia (*Oreochromis niloticus*) en Malpaso, Chiapas. *Espacio I+D*, Innovación más Desarrollo, 10(27). https://espacioimasd.unach.mx/index.php/Inicio/article/view/270/875#:~:text=La%20temperatura%20promedio%20encontrada%20en,la%20incidencia%20de%20la%20bacteria.

INEGI (2023) www.inegi.org.mx [Accessed April 2023].

Kayansamruaj, P., Pirarat, N., Katagiri, T., Hirono, I., & Rodkhum, C. (2014). Molecular characterization and virulence gene profiling of pathogenic *Streptococcus agalactiae* populations from tilapia (*Oreochromis* sp.) farms in Thailand. Journal of Veterinary Diagnostic Investigation, 26(4), 488– 495. https://doi.org/10.1177/1040638714534237

Lau, S. K. P., Woo, P. C. Y., Tse, H., Leung, K.W., Wong, S. S. Y. & Yuen, K.Y. (2003). Invasive *Streptococcus iniae* infections outside North America. Journal of Clinical Microbiology, 41, 1004–1009. https://doi.org/10.1128/JCM.41.3.1004-1009.2003

Lebel, L., Lebel, P. & Lebel, B. (2016). Impacts, perceptions and management of climate-related risks to cage aquaculture in the reservoirs of Northern Thailand. Environmental Management, 58(6), 931–945. https://doi.org/10.1007/s00267-016-0764-5

Li, Y., & Cai, S.H. (2011). Identification and pathogenicity of *Aeromonas sobria* on tail rot disease in juvenile tilapia *Oreochromis niloticus*. Current Microbiology, 62(2), 623– 627. https://doi.org/10.1007/s00284-010-9753-8

Li, Y. W., Liu, L., Huang, P. R., Fang, W., Luo, Z. P., Peng, H. L., Wang, Y. X. & Li, A. X. (2014). Chronic streptococcosis in Nile tilapia, *Oreochromis niloticus* (L.), caused by *Streptococcus agalactiae*. Journal of Fish Diseases, 37(8), 757–763. https://doi.org/10.1111/jfd.12146

Liu, Y., Li, L., Huang, T., Wu, W., Liang, W., & Chen, M. (2019). The interaction between phagocytes and *Streptococcus agalactiae* (GBS) mediated by the activated complement system is the key to GBS inducing acute bacterial meningitis of tilapia. Animals, 9(10), 818. https://doi.org/10.3390/ani9100818

Martínez-Cordero, F. J., Delgadillo, T. S., Sanchez-Zazueta, E., & Cai, J. (2021). Tilapia aquaculture in Mexico-assessment with a focus on social and economic performance. FAO, Rome, No. 1219. https://doi.org/10.4060/cb3290en

Martínez-Murcia, A. J., Monera, A., Saavedra, M. J., Oncina, R., Lopez-Alvarez, M., Lara, E., & Figueras, M. J. (2011). Multilocus phylogenetic analysis of the genus *Aeromonas*. Systematic and Applied Microbiology, 34(3), 189–199. https://doi.org/10.1016/j.syapm.2010.11.014

Maulu, S., Hasimuna, O. J., Mphande, J., & Munang’andu, H. M. (2021). Prevention and control of streptococcosis in tilapia culture: a systematic review. Journal of Aquatic Animal Health, 33(3), 162–177. https://doi.org/10.1111/jfd.12146

Mian, G. F., Godoy, D.T., Leal, C.A., Yuhara, T.Y., Costa, G.M. & Figueiredo, H.C. (2009). Aspects of the natural history and virulence of *Streptococcus agalactiae* infection in Nile tilapia. Veterinary Microbiology, 36(1–2), 180–183. https://doi.org/10.1016/j.vetmic.2008.10.016

OIE (2022). https://www.woah.org/en/what-we-do/standards/codes-and-manuals/aquatic-code-online-access/?id=169&L=1&htmfile=chapitre_welfare_stunning_killing.htm [Accessed June 23 2023]

Pridgeon, J. W., & Klesius, P. H. (2011). Virulence of *Aeromonas hydrophila* to channel catfish *Ictaluras punctatus* fingerlings in the presence and absence of bacterial extracellular products. Diseases of Aquatic Organisms, 95(3), 209–215. https://doi.org/10.3354/dao02357

Ortega, C., García, I., Irgang, R., Fajardo, R., Tapia-Cammas, D., Acosta, J., & Avendaño-Herrera, R. (2018). First identification and characterization of *Streptococcus iniae* obtained from tilapia (*Oreochromis aureus*) farmed in Mexico. Journal of Fish Diseases, 41(5), 773–782. https://doi.org/10.1111/jfd.12775

Rahmatullah, M., Ariff, M., Kahieshesfandiari, M., Daud, H. M., Zamri-Saad, M., Sabri, M. Y., Amal, M. N. A., & Ina-Salwany, M. Y. (2017). Isolation and pathogenicity of *Streptococcus iniae* in cultured red hybrid tilapia in Malaysia. Journal of Aquatic Animal Health, 29(4), 208–213. https://doi.org/10.1080/08997659.2017.1360411

SAGARPA (2022). Against streptococcosis … good aquaculture practices. México. (Last view March 2023). https://www.gob.mx/agricultura/articulos/contra-la-estreptococcosis-buenas-practicas-acuicolas [Accessed 20 June 2023]

Soliman, M., Ellakany, H., Gaafar, A., Elbialy, A., Zaki, M., Younes, A. (2014). Epidemiology and antimicrobial activity of methicillin-resistant *Staphylococcus aureus* (MRSA) isolated from Nile tilapia (*Oreochromis niloticus*) during an outbreak in Egypt. Life Science Journal, 11, 1245–1252

Soto, E., Griffin, M., Arauz, M., Riofrio, A., Martinez, A., & Cabrejos, M. E. (2012). *Edwardsiella ictaluri* as the causative agent of mortality in cultured Nile tilapia. Journal of Aquatic Animal Health, 24(2), 81– 90. https://doi.org/10.1080/08997659.2012.675931

Soto, E., Kidd, S., Mendez, S., Marancik, D., Revan, F., Hiltchie, D., & Camus, A. (2013). *Francisella noatunensis* subsp. *orientalis* pathogenesis analyzed by experimental immersion challenge in Nile tilapia, *Oreochromis niloticus* (L.). Veterinary Microbiology, 164(1-2), 77–84. https://doi.org/10.1016/j.vetmic.2013.01.024

Soto-Rodriguez, S. A. (2009). Calidad del agua y bacterias presenetes en tilapia cultivada (Water quality and bacteria in cultured tilapia). Mexico. https://www.fps.org.mx/portal/index.php/publicaciones/99-acuicolas/1088-calidad-del-agua-y-bacterias-presentes-en-tilapia-cultivada [Accessed June 20 2023]

Soto-Rodriguez, S. A., Cabanillas-Ramos, J., Alcaraz, U., Gomez-Gil, B., & Romalde, J. L. (2013). Identification and virulence of *Aeromonas dhakensis*, *Pseudomonas mosselii* and *Microbacterium paraoxydans* isolated from Nile tilapia, *Oreochromis niloticus*, cultivated in Mexico. Journal of Applied Microbiology, 115(3), 654–662. https://doi.org/10.1111/jam.12280

Soto-Rodriguez, S. A., Lozano-Olvera, R., Garcia-Gasca, M.T., Abad-Rosales, S.M., Gomez-Gil, B. & Ayala-Arellano, J. (2018). Virulence of the fish pathogen *Aeromonas dhakensis*: genes involved, characterization and histopathology of experimentally infected hybrid tilapia. Diseases of Aquatic Organisms, 129(2), 107–116. https://doi.org/10.3354/dao03247

Watanabe, W. O., Losordo, T. M., Fitzsimmons, K., & Hanley, F. (2002). Tilapia production systems in the Americas: technological advances, trends, and challenges. Reviews in Fisheries Science, 10(3-4), 465–498. https://doi.org/10.1080/20026491051758

Zhou, S. M., Fan, Y., Zhu, X. Q., Xie, M. Q., & Li, A. X. (2011). Rapid identification of *Streptococcus iniae* by specific PCR assay utilizing genetic markers in ITS rDNA. Journal of Fish Diseases, 34(4), 265–271. https://doi.org/10.1111/j.1365-2761.2010.01233.x

