## Supplementary material for "Prevalence of pathogenic bacteria detected by qPCR from cultured Nile tilapia (*Oreochromis niloticus* Linnaeus, 1758) in southwest Mexico"

**Short running title:** pathogenic bacteria from farmed tilapia

**Keywords**: fish pathogens, qPCR, cultured Nile tilapia, southwest Mexico

**a**

**1,100 bp**

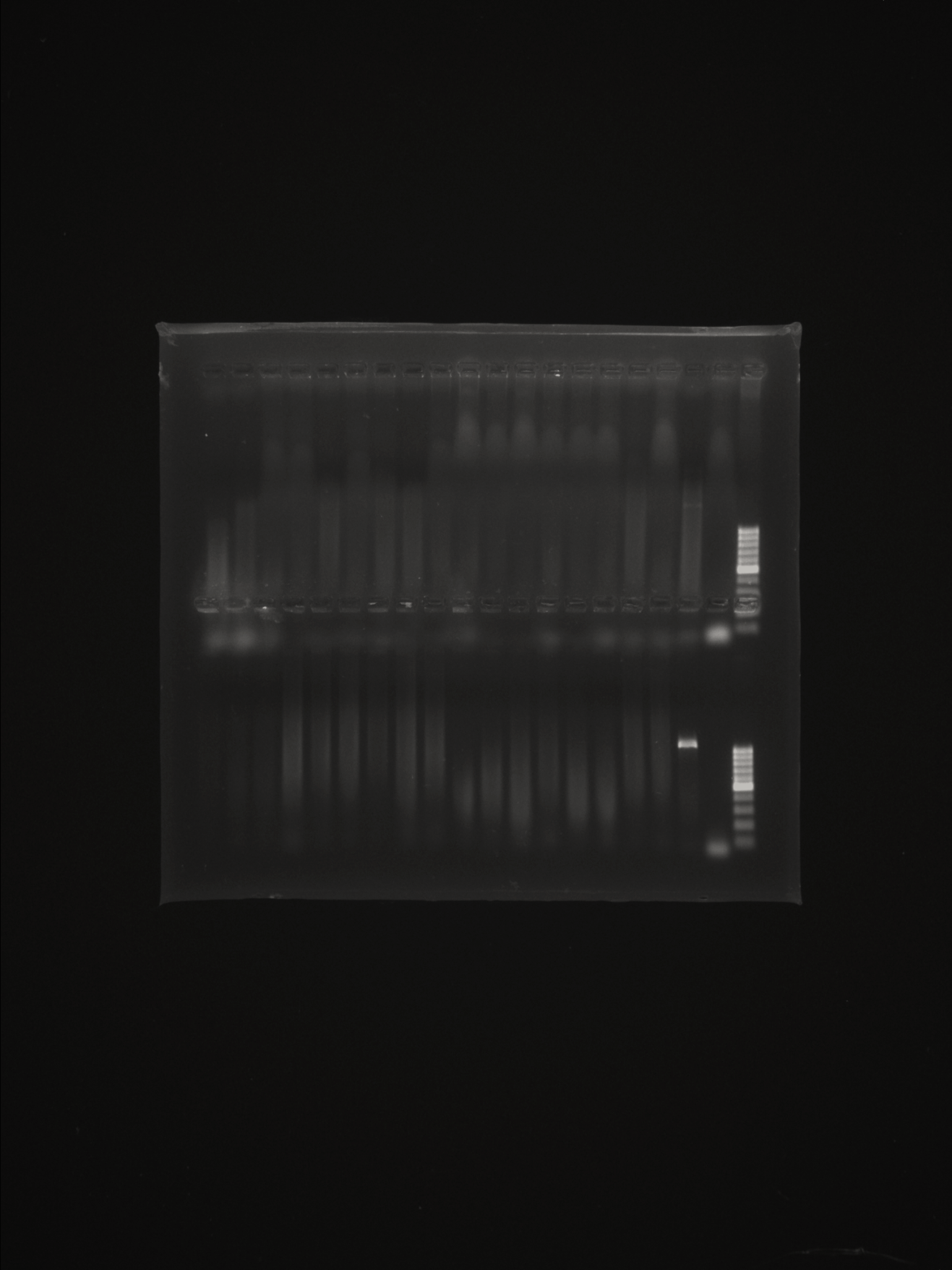

**B**

**b**

**a**

**a**

Fig S1. Amplification of *Francisella noatunensis* PCR products. **A.** line a, molecular marker; line b, negative control; lines c and d, positive control of *F. noatunensis* sub*. orientalis*. **B**, line a, molecular marker; b, negative control; rest of the lines organ tissue samples of *O. niloticus* from Chiapas in 2019 without amplification.

| Table S1. Water physicochemical parameters taken in the sampling site of tilapia for year and state. | | | | |
| --- | --- | --- | --- | --- |
| Year | State | Temperature (°C) | Dissolved oxygen (mg/L) | pH |
| 2018 | Guerrero | 26-32 | 6.5-11.0 | 8.1-9.0 |
|  | Oaxaca | 21-27 | 2.8-10.5 | 7.5-9.5 |
|  | Chiapas | 22-30 | 7.9-1.0 | 7.1-8.2 |
| 2019 | Guerrero | 30-32 | 2.3-7.7 | 7.8-8.8 |
|  | Oaxaca | 24-32 | 4.1-8.6 | 7.0 |
|  | Chiapas | 26-33 | 3.5-13.0 | 7.4-7.6 |

1. * Correspondence: Sonia Araceli Soto-Rodriguez, Centro de Investigación en Alimentación y Desarrollo, Unidad Mazatlán en Acuicultura y Manejo Ambiental, Av. Sábalo Cerritos, Mazatlán, Sinaloa, México.
